# A harmonised systems-wide re-analysis of greenhouse gas emissions from sunflower oil production

**DOI:** 10.1101/2020.06.19.161893

**Authors:** Thomas D Alcock, David E Salt, Stephen J Ramsden

## Abstract

Sunflower (*Helianthus annuus* L.) is the largest source of vegetable oil in Europe and the fourth largest globally. Intensive cultivation and post-harvest steps contribute to global food-systems’ greenhouse gas (GHG) emissions. However, variation between production systems and reporting disparity have resulted in discordance in previous emissions estimates. To assess systems-wide GHG implications of meeting increasing edible oil demand using sunflower, we performed a unified re-analysis of primary life cycle inventory data, representing 995 farms in 11 countries, from a saturating search of published literature. Total GHG emissions varied from 1.1 to 4.2 kg CO_2_-equivalent per kg oil across systems, 62% of which originated from cultivation. Major emissions sources included diesel- and fertiliser-use, with irrigation electricity contributing most to between-systems variation. Our harmonised, cross-study re-analysis not only enabled robust comparisons and identification of mitigation opportunities across sunflower oil production systems, but also lays the groundwork for comparisons between alternative oil crops.

## Introduction

From around 800,000 years before the common era, up to the year 1800, atmospheric carbon dioxide (CO_2_) concentrations averaged 228 parts per million (ppm)^1^. Despite regular fluctuations, coinciding with ice ages and interglacial periods, concentrations never rose above 300 ppm during this time. However, since the industrial revolution, atmospheric CO_2_ has been rising year on year^2^. Since 1913, concentrations have failed to drop below 300 ppm. In every year since 2015, they have remained above 400 ppm, 70-80 % higher than pre-industrial concentrations^3, 4^. The Intergovernmental Panel on Climate Change (IPCC) stated that the dominant cause of global warming since 1950 has been anthropogenic contributions to greenhouse gas (GHG) emissions^5^. As the human population has grown, food production has risen markedly; today, food supply chains are responsible for 26% of all GHG emissions^6^. As we strive to provide greater amounts of nutritious food to over 800 million currently undernourished people^7^, whilst meeting additional demand as the population continues to grow^8^, carefully targeted global food system interventions will form a key part of our efforts to limit the effects this has on planetary health.

Vegetable oils are a major source of dietary polyunsaturated fatty acids^9^ and are a crucial component of a wide range of cuisines. Steadily increasing demand for edible oil has led to increased oil crop production^10^ through expansion of cultivation area^11^, and intensification of production practices^12^. Oil crops account for over 300 million hectares globally, approximately 19% of total cropped land (excluding pasture)^10^. Thus, they represent a major and growing source of GHG emissions^13^. Sunflower is the largest source of vegetable oil in Europe, and fourth largest globally, with over 26 million hectares harvested per year^10^. Estimates of GHG emissions produced by sunflower production have been published previously, many of which follow life cycle assessment (LCA) methodology^14^. Significant variation exists between assessment results, much of which is due to regional variation and different production practices. For example, life cycle GHG emissions, measured as CO_2_-equivalent (CO_2_e), from sunflower seed production in Italy were estimated to be approximately 0.6 kg CO_2_e per kg seed^15^, whilst in Iran, where large quantities of irrigation water are required for crop growth, emissions were estimated as high as 2.3 kg CO_2_e per kg seed^16^. However, despite genuine differences, it is likely that much of the variation between studies is a result of non-harmonious reporting. Additionally, functional units often vary wildly between studies, further complicating meaningful comparison.

This work aimed to determine system-specific GHG implications arising from meeting additional, future demand for vegetable oil using sunflower. This was achieved using a meta-analytical approach, whereby production inventory data from all relevant literature published over the last 20 years was isolated and used to re-calculate methodologically consistent life cycle GHG emissions estimates. This analysis will help to identify systems capable of meeting future demand as sustainably as possible, whilst laying the groundwork for future comparison of emissions from sunflower oil production with those associated with other major oil crop systems such as palm, soybean and rapeseed.

## Methods

### Study objective and strategy

The aim of this study was to critically review global variation in sunflower oil life cycle greenhouse gas (GHG) emissions, as part of a wider project aiming to address the sustainability of the world’s major oil crops. The approach taken was that of a comprehensive re-analysis of primary data sources, whereby relevant literature was systematically isolated from a saturating search of multiple bibliographic databases, data extracted, normalised, and used to conduct novel comparisons. Specifically, raw energy and material input data was collected from studies that followed life cycle assessment (LCA) methodology^14^ and used to curate an inventory of inputs required for sunflower oil production. Extraction of raw energy and material data from the literature, rather than reported emissions values, was crucial for achieving consistent and comparable results, due to variation in emissions calculations, system boundaries, and functional and time units between studies. Where possible, individual studies were grouped into distinct production systems based on geographic range, cultivation and processing practices, in order to reduce within-study error. Inventory data were then used to calculate associated life cycle GHG emissions based on a series of conversion factors, following a consequential LCA modelling approach.

### Consequential Modelling Approach

A consequential LCA models the effects that production and use of a given product have on environmental burdens. This differs from an attributional LCA, which instead models the proportion of current environmental burdens that belong to the studied product^17^. The definitions are seemingly nuanced, but the methodology chosen has profound implications for the study results. This is particularly relevant if a process being studied produces multiple, inseparable products, as is the case for sunflower cultivation and processing, which produces both sunflower oil and sunflower cake, the latter used as protein-rich animal feed. In *attributional* LCA, total environmental burdens of such processes are proportionately allocated to each co-product based on mass, economic value, or other comparative metrics^18^. This allocation of environmental burden to different products is problematic when attempting to assess the impact of change in any one co-product: e.g. in the context here, the consequences of an increase in sunflower production are not just a consequence of the oil component, as there will also be an increase in cake for animal feed. In contrast, *consequential* LCA assumes that a decision to buy or use a product drives demand for it, which drives a larger, ‘joint’ production process, especially if the product under study is deemed the main product in the joint system^19^. Crucially, the implication is that the product is produced by additional capacity and thus the production system is expanded to include emissions produced by the co-products as well^20^. An objective of this study was to evaluate the GHG implications of increased sunflower oil production by comparison of the environmental burden of production in different locations and using different practices. For this reason, and since sunflower oil and cake production in each system are inextricably linked, consequential LCA modelling was selected and used throughout. This is likely to be of much greater relevance to decision making, and for future comparison of the impacts of meeting increasing demand for vegetable oil between different crop systems.

### System boundaries

Sunflower production systems in the meta-analysis were modelled from cradle to factory-gate. Each system studied was split into five production stages: 1. cultivation and harvest, 2. seed drying and storage, 3. transport to processing facilities, 4. processing and refining, and 5. packaging. These were determined based on stages reported in studies included in our meta-analysis. Production stages are summarised in Figure 1, in which major inputs and sources of direct emissions are noted. Land-use change was omitted from the model, on the basis that global expansion of land used to cultivate sunflower has slowed in recent decades to only around 1-2 % per year on average (FAO, 2020). Stages post-packaging such as distribution and use are also omitted, since these are more driven by consumer preferences than decisions made during production. Within the cultivation stage, material inputs comprised diesel and lubricants required to run agricultural machinery, seeds for planting, fertilisers, manure, pest and disease controls, and steel used in agricultural machinery, as a proportion of its useful life. Electricity required to power irrigation (where present) was further included as an energy input. The second stage comprised electricity and biomass used for combustion to reduce seed moisture concentrations ready for processing and for storage prior to transfer to processing facilities. The transport stage reports GHG emissions produced during transport of seed to processing facilities. This was measured in tonne kilometres (tkm), which represents the transport of one tonne of goods over a distance on one kilometre via a given mode of transit. Processing and refining comprise electricity, fuel and lubricants required for seed pressing, steam and chemical inputs for further oil extraction and refining, and metals, concrete and asphalt used in the processing facility, as a proportion of their useful life. Finally, the packaging stage reports all plastics used to produce oil bottles, as well as plastics, cardboard and wooden pallets used to further package the finished product ready for distribution. Inputs of services such as cleaning, marketing, accounting, as well as overheads including office space electricity and upkeep were omitted from the model, due to a lack of reporting in studies included in the meta-analysis.

**Figure 1.**
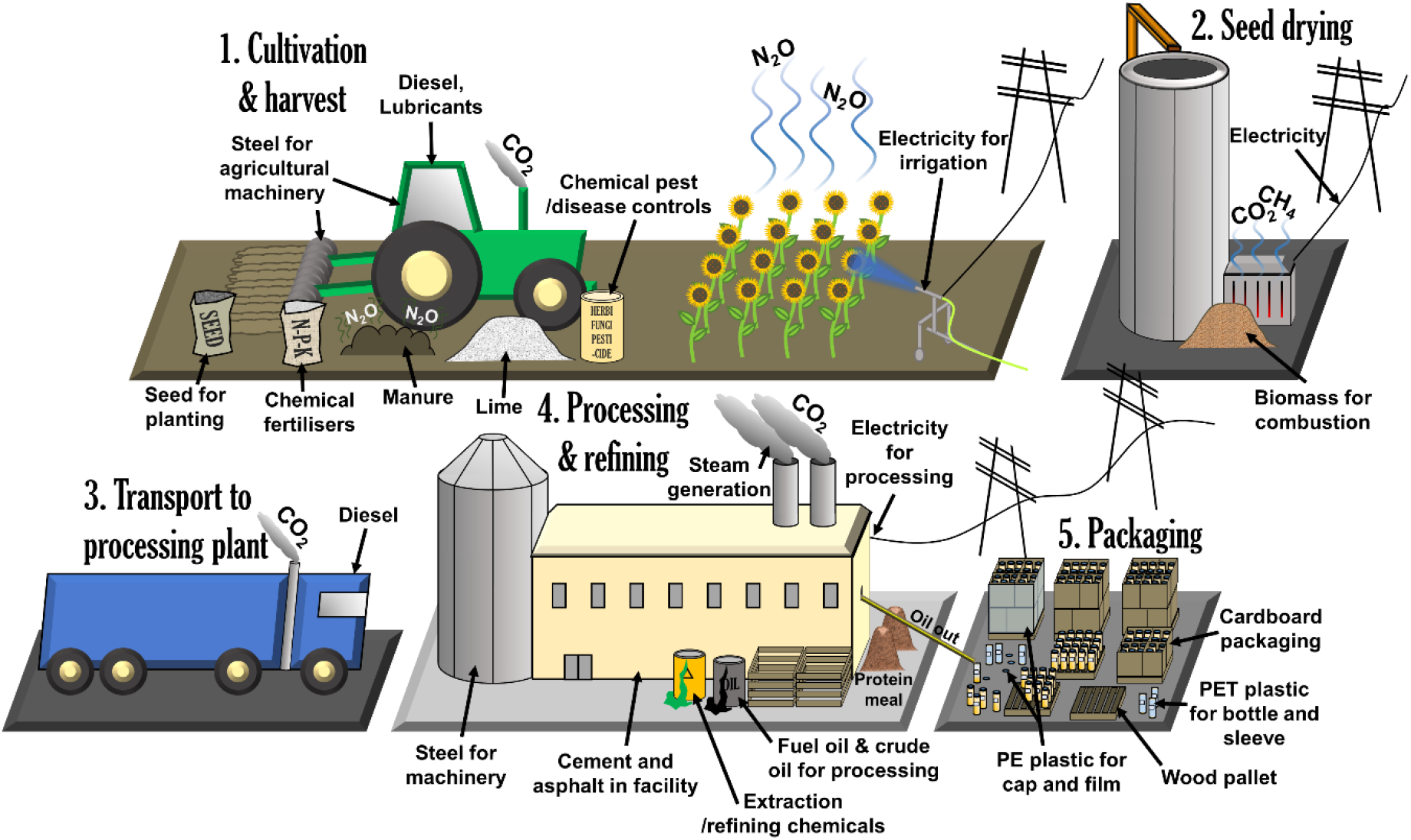
System boundaries of life cycle assessment model (not to scale). Emissions sources and major direct emissions shown.

### Functional units

Two major functional units are referred to throughout this study. For systems modelling and spreadsheet management, energy and material inputs are referred to on a per hectare (ha) basis, since this unit is most relevant to decision making at the cultivation stage. For the purpose of final results reporting, as well as management of the packaging stage of the systems under study, the functional unit is defined as one kg of refined, packaged sunflower oil, which best reflects the scope of this work.

### Literature eligibility criteria

Studies were assessed for eligibility for inclusion using nine criteria (Table 1). These were formulated to fulfil the PRISMA statement reporting guidelines, designed to promote transparent and complete reporting of systematic reviews and meta-analyses^21^. Critically, literature was only included in this study if it was published in English over the last 20 years, from the year 2000 (inclusive) up to the early part of 2020. Literature was also required to significantly concern production of sunflower over other crops, specifically in a setting that the present authors deemed to be commercially viable as opposed to experimental or speculative (e.g. on abandoned quarries) and in the context of sustainability. To ensure studies included data relevant to the aims of our meta-analysis, inclusion criteria #7 through #9 were also introduced. This enabled isolation of studies which contained LCA-style inventory data whilst omitting studies that didn’t contain data relevant to the system boundaries under study here. Studies were also required to frame their inventory data in terms of one or more of the functional units used here or enable recalculation into such units based on available data.

**Table 1.** Inclusion criteria for studies to be used in the meta-analysis.

| # | Study inclusion criteria |
| --- | --- |
| 1 | Published in the English language |
| 2 | Original, unique research article, conference paper, or PhD thesis with clear study methodology |
| 3 | Published between the years 2000 and 2020 (inclusive) |
| 4 | Significantly concerns production of sunflower (*Helianthus annuus*) seed as a source of edible vegetable oil. Studies concerning use for biofuel production also included, providing data from relevant earlier stages of production were reported |
| 5 | Concerns production in a relevant, commercially viable setting i.e. not experimental/speculative |
| 6 | Concerns sustainability in the context of greenhouse gas emissions |
| 7 | Uses Life Cycle Assessment or similar methodology |
| 8 | Contains inventory data relevant to our system boundaries (**Figure 1**), sufficient to calculate associated greenhouse gas emissions |
| 9 | Presents data in terms of our functional units, or enables recalculation using reported inventory data |

### Information sources, search strings and record compilation

In order to thoroughly extract all relevant literature, eight individual bibliographic databases were consulted. These were Web of Science (all databases), Scopus, PubMed, PubMed Central, Wiley Online Library, SpringerLink, JSTOR, and ScienceDirect. These databases were selected based on their multidisciplinary content, search string capacity and overall performance, as analysed by Gusenbauer and Haddaway^22^. Search strings were formulated to identify studies that concerned sunflower in the context of oil production and sustainability and/or involved LCA. Biofuel/biodiesel was also included in the search strings to incorporate studies which may include data relating to sunflower cultivation and relevant processing steps without specifically referring to them. Search strings varied depending on the required syntax of each bibliographic database, but broadly followed the string used for Web of Science as per below:

[(“sunflower” OR “helianthus”) (“life cycle assessment” OR “life cycle analysis” OR “lca” OR “greenhouse gas emissions” OR “greenhouse emissions” OR “carbon footprint” OR “sequestration” OR “nutrient loss”) (“oil” OR “biodiesel” OR “biofuel”)]

Full search strings used for all other databases are included in Supplementary Table 1 along with number of search results returned for each. An additional search was carried out in Web of Science filtered to only include results from *The International Journal of Life Cycle Assessment* with the search string [“sunflower” OR “helianthus”] to reduce the risk of missing highly relevant literature that wasn’t identified with the main search strings. In general, and where possible for each database, searches were directed to scan only text in the title, abstract and in any keywords, since searching in full text records led to too many spurious results. Searches were performed on 13^th^ February 2020.

All records identified through the above methods were downloaded and imported into EndNote X9 (Clarivate Analytics, Philadelphia, PA, USA). Where databases didn’t allow export of records in a format recognised by EndNote, they were exported as text files and converted to EndNote readable formats in Microsoft Excel.

### Screening

Since multiple bibliographic databases were used to identify relevant literature, many records were represented more than once in the initial EndNote library. This was addressed first by sorting records alphabetically and manually removing duplicates, then by using the EndNote “Find Duplicates” tool to address any that were missed. All remaining, unique records were then exported using custom output styles to Excel for screening. Records were initially screened based on publication year, language and type, then by titles and finally abstract, in order to quickly exclude literature which obviously failed to fulfil the inclusion criteria (Table 1). Full text articles were accessed online for the remaining records. Text, tables and figures, and supplementary information was consulted to ensure that only relevant literature was retained for analysis. On occasion, unique records corresponding to the same study were identified, for example where a conference paper was submitted prior to a full journal submission. In these cases, only the most complete or recent record was retained.

### Data collection process

Data collection for the meta-analysis utilised a custom inventory spreadsheet managed in Microsoft Excel. Each literature record was given a unique source identifier and allocated to a unique row within the spreadsheet. Summary information including study location, soil type, cultivation and oil extraction method was noted. Records were then accessed in turn and data was identified in tables, figures, text and supplementary information (where available). All data relevant to the system boundaries of the current study were extracted and used to populate the inventory spreadsheet. The reporting of certain data items was simplified in the inventory to provide a suitable number of values for comparison. For example, chemical disease/pest controls were grouped into herbicide, insecticide, fungicide and unspecified pesticide items, rather than reporting specific chemicals used. Similarly, fertilisers were grouped into major data items including mineral nitrogen (N), urea N, manure (total weight), phosphate (as P_2_O_5_), and potassium oxide (K_2_O). Inventory data were all expressed in terms of the functional unit “per hectare”. Data that were expressed in alternative units in the literature were converted using other available data. For example, where inventory data were expressed in terms of “per kg refined oil”, values were adjusted to “per hectare” by dividing by reported oil recovery as a proportion of 1, then multiplying by reported seed yield in kg per hectare. Study-specific inventory data were used to perform conversions as much as possible. However, values were assumed in cases where such information was not available, including from average values reported in other relevant literature in the inventory spreadsheet and in online databases including FAOSTAT^10^. For the sake of simplicity, consistent units per hectare were utilised for individual energy and material data items, including kg for diesel, lubricants, fertilisers, liquid disease control, steel for machinery and other physical categories, and MJ for energy-based categories. Where these were reported differently in each literature record, values were converted using consistent conversion ratios e.g. 1 kWh = 3.6 MJ, 1 L diesel = 0.832 kg. A full list of data items collected, conversion factors used, and assumptions made are reported in Supplementary Table 2.

### Assessing risk of bias and record consolidation

It was assumed that reporting bias existed within studies, including variation in sources of inventory data, choice of inventory items, system boundaries, and choice of using actual on-farm/system data, survey data, regional average data or unverified assumptions. Bias was also assumed across studies, including underrepresentation of some systems in the literature. In order to highlight and where possible address this, the following measures were taken. For each record, it was noted what kind of system was used to acquire inventory data. Where this was survey data, the number of survey participants was noted; where this was production data, the number of farms represented by the data was noted. Records were then consolidated into several specific production systems, based on geographic production range and cultivation/processing methods, as per Poore and Nemecek^6^. For each data item for each system, the mean of all reported values was then calculated and used as the system standardised value. Where data items relevant to the system boundaries of this study were not reported in individual literature records, cells were generally left blank in the inventory database. The exception was where it was deemed likely that the true value for a specified category was zero if not reported. For example, if a study reported kg of urea N applied to a field but failed to mention mineral N, it was assumed that no mineral N was used. In these cases, zero values were added to relevant cells in the inventory. Thus, blank cells were left out of subsequent analyses, and imputed zero values were included, on the assumption that they were representative of within-system variation. Decisions on whether to leave cells blank or impute zero values for each data item are summarised in Supplementary Table 2. For each system, the number of studies combined and the collective number of farms/survey participants that each represented was noted. This approach enabled the majority of data categories to be filled for each system. Where systems were still missing a value for a given data item, the mean average value of data item values across all systems was calculated and used. Where appropriate, this was weighted by system yield. This was most relevant for steps post-cultivation, e.g. chemicals used for processing and refining sunflower seeds, which are a product of quantity of seed processed rather than area of land cultivated. Finally, the number of records present for each system was compared to global, country-specific production data from FAOSTAT^10^, to identify any disparities between production quantity and scientific reporting incidence.

### Formulation of GHG emission factors database and emissions reporting

To enable calculation of greenhouse gas emissions from the inventory data, an emission factors database was compiled. This consisted of estimated carbon dioxide (CO_2_), methane (CH_4_) and nitrous oxide (N_2_O) emissions associated with the manufacture, distribution and use of the energy and material inputs under study here. The collection of emission factors relating to the three gasses individually allowed consistent calculation of CO_2_-equivalent (CO_2_e) emission factors, which form an effective means to quantify Global Warming Potential (GWP). For this study, we used IPCC AR5 GWP_100_ conversion factors with climate-carbon feedbacks^23^. Emission factors were collected from multiple emissions databases including BioGrace^24^, UK Government GHG Conversion Factors for Company Reporting 2019^25^ the EMEP/EEA air pollutant emission inventory guidebook 2019^26^ and the software GREET 2019 (version 1.3, Argonne National Laboratory, IL, USA), or from literature sources. Electricity emissions were calculated using country-specific emission factors to reflect regional variation in electricity generation practices. These were taken from the BioGrace Additional Standard Values database, based on International Energy Agency (Paris, France) data from 2012, a year which is roughly central in our study period. For some data items, only CO_2_e emission factor values were available, many of which were calculated using previous GWP conversion estimates. Where recalculation to AR5 values wasn’t possible, these were retained as a best estimate of the emissions associated with the given factor. It is worthy of note that, of the gasses under study here, only the CH_4_ conversion factor differs between IPCC AR4 and AR5 (with climate-carbon feedbacks). Hence, for data items for which AR4 conversion factors are used here, it is likely that only minimal error in final emissions calculations exists.

### Summary measures and statistical analysis

Once emissions values for each data item for each system were calculated, the sum of emissions values for each production stage (cultivation, transport, seed drying, processing, packaging), and total life cycle emissions were determined and reported as global warming potential (GWP; CO_2_-equivalent). Comparisons were drawn between emissions from the different data items to ascertain those which contributed the most to total life cycle emissions. Violin plots were generated in GraphPad Prism 8 (GraphPad Software, San Diego, CA, USA) for visualisation of within-production stage variation, with mean, median and upper and lower quartiles for each stage shown. Further summary statistics including correlation and regression analyses between data items and emissions were generated in GenStat (20^th^ ed. VSN International, Hemel Hempstead, UK). Regression plots were generated in GraphPad Prism 8.

## Results

### Inventory data obtained for 22 distinct production systems, with gaps in the literature revealed

Inventory data for inclusion in the meta-analysis were obtained via a comprehensive literature search strategy involving eight separate bibliographic databases. The initial search identified 408 records, which were reduced to 271 unique records once duplicates from the different databases were removed (Figure 2). Records were assessed based on the criteria listed in Table 1. Firstly, by screening the title and abstract, 136 records were excluded, leaving 135. Full text articles and associated data were then accessed, from which a further 103 records were excluded. This resulted in the isolation of 32 articles which contained data relevant to our inclusion criteria, representing production in 11 countries. These are included as additional references in Supplementary Table 3. Fifteen of those records (47%) originated from Italy, which formed the single largest regional data source (Figure 3a). Within all 32 records, 58 production systems were reported (many records investigated more than one system). These comprised inventory data representative of 995 farms (Table 2), either through direct observation or survey, as well as additional inventory data previously reported in the literature or hypothetically modelled. Thirty of these reports (52%) specifically concerned sunflower production in Italy, with other countries represented to lesser degrees (Figure 3b).

**Figure 2.**
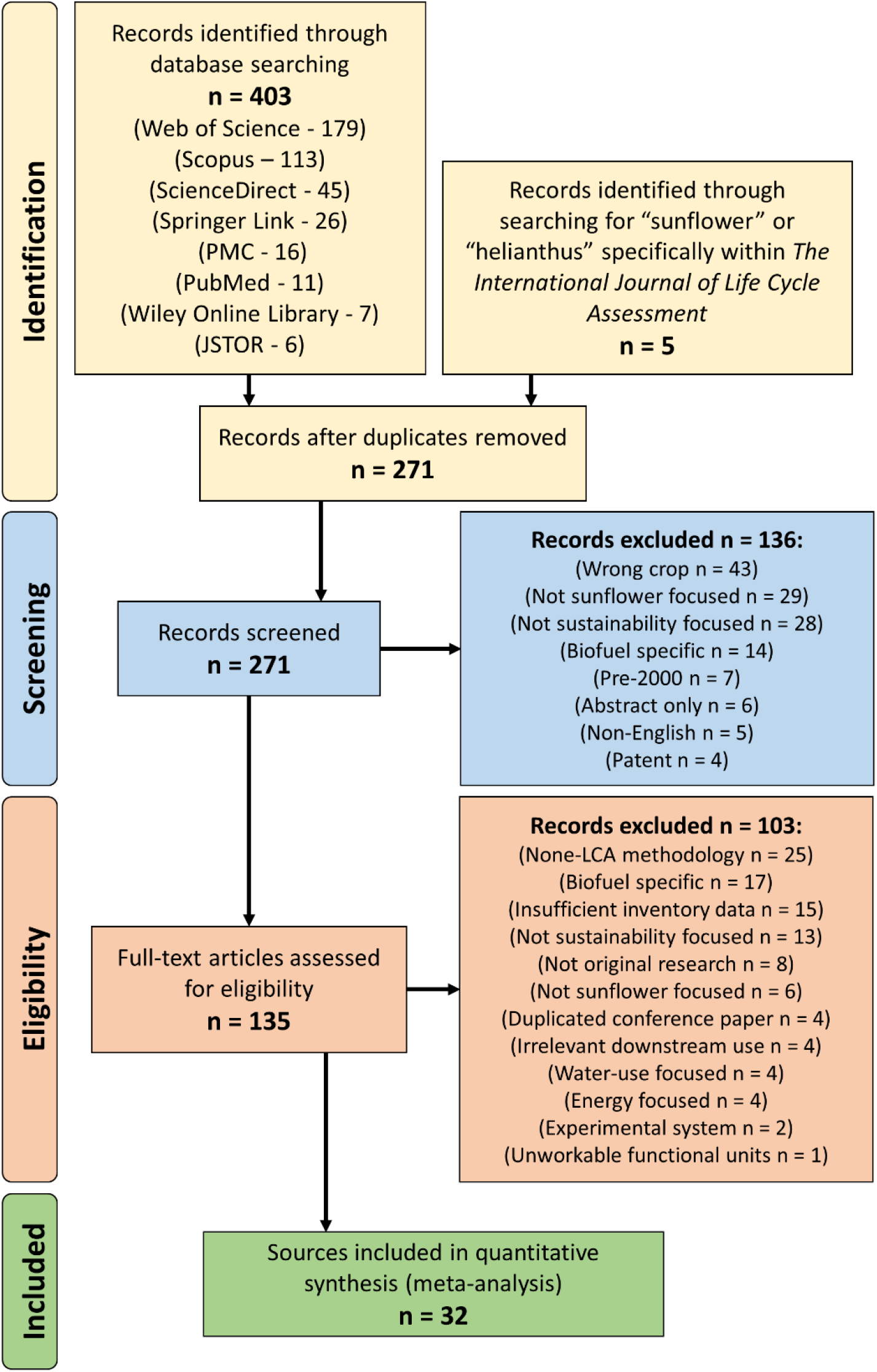
PRISMA flow diagram, showing number of records identified from literature searching, and reasons for exclusion. Note some records failed to meet multiple inclusion criteria. In these cases, the principle reason for exclusion is listed. Hence, each record only appears in this figure once.

**Figure 3.**
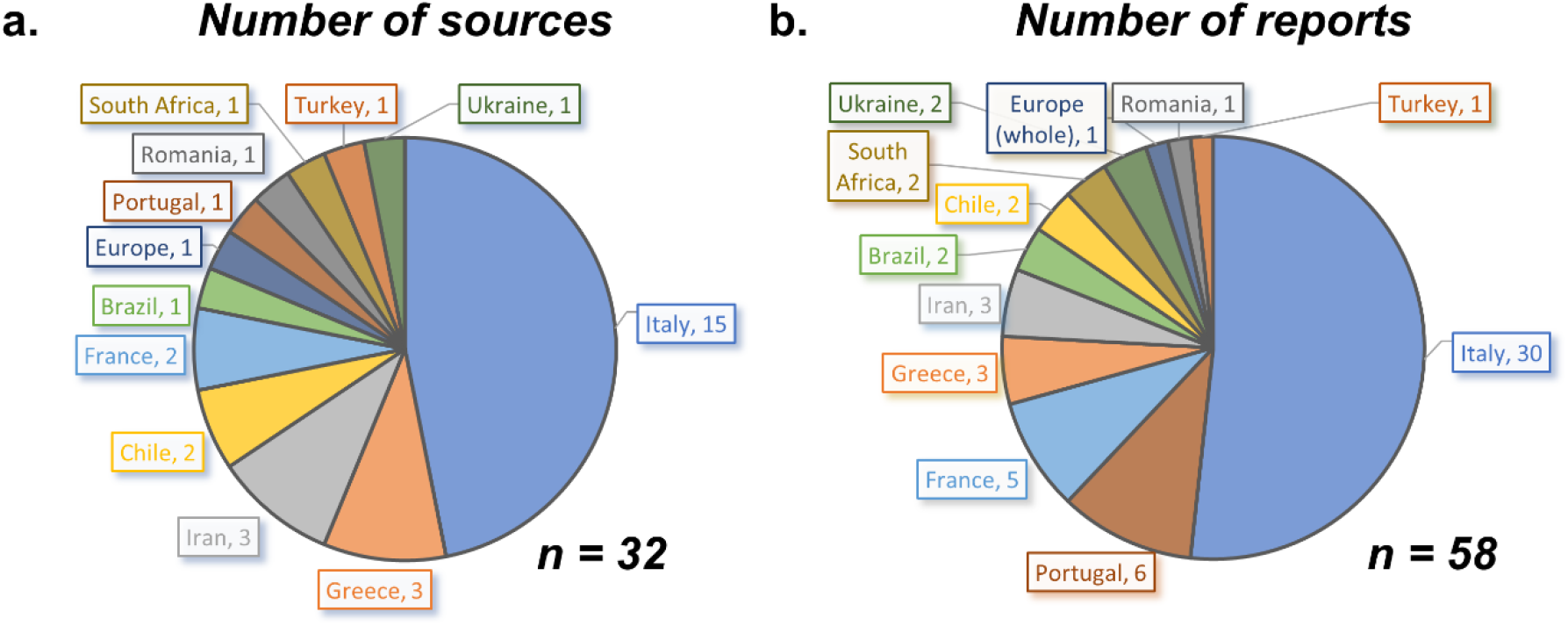
Study locations of sources (a.) and reports (b.) included in the meta-analysis. ‘Sources’ refers to individual, unique articles/records in the literature, whereas ‘reports’ refers to individual sunflower production systems within sources.

**Table 2.** Consolidated systems reporting used in meta-analysis, number of reports included per system, and, where relevant, number of farms represented based on observation or survey (“N/A” = data based only on hypothetical or literature values). System yield and oil recovery included for comparison (“-” = data not reported). GWP (CO2equivalent) reported on a per hectare (ha) and per kg oil basis.

| System name | No. Reports | Farms represented | Yield (t ha<sup>-1</sup>) | Oil recovery (% w/w) | GWP per ha | GWP per kg oil |
| --- | --- | --- | --- | --- | --- | --- |
| Brazil, conventional | 1 | 5 | 1774 | - | 970 | 1.40 |
| Brazil, monoculture | 1 | N/A | 1774 | - | 1013 | 1.47 |
| Chile, conventional | 2 | N/A | 2200 | 37 | 1425 | 1.75 |
| Italy, conventional | 27 | 164 | 2283 | 41 | 1929 | 2.06 |
| Italy, low intensive | 1 | N/A | 2500 | - | 1301 | 1.33 |
| Italy, cold press, conventional | 1 | 5 | 2000 | 33 | 1591 | 2.41 |
| Italy, cold press, organic | 1 | 5 | 1400 | 33 | 592 | 1.28 |
| France conventional | 1 | N/A | 2169 | - | 1243 | 1.47 |
| France, no legume, bare | 1 | 1 | 2678 | - | 1429 | 1.37 |
| France, no legume, cover crop | 1 | 1 | 2506 | - | 1395 | 1.43 |
| France, legume, bare | 1 | 1 | 2581 | - | 1116 | 1.11 |
| France, legume, cover crop | 1 | 1 | 2673 | - | 1139 | 1.08 |
| Portugal, irrigated | 3 | ~400 | 3287 | - | 1862 | 1.45 |
| Portugal, rainfed | 3 | ~400 (as above) | 943 | - | 697 | 1.90 |
| Greece, conventional | 3 | 6 | 2563 | 36.5 | 1781 | 1.90 |
| Iran, conventional | 3 | 73 | 2339 | - | 3822 | 4.19 |
| Romania, conventional | 1 | N/A | 1210 | - | 1135 | 2.41 |
| South Africa, conventional | 1 | 1 | 950 | - | 917 | 2.48 |
| South Africa, recommended | 1 | N/A | 2000 | - | 1642 | 2.11 |
| Turkey, conventional | 1 | 332 | 1870 | - | 1295 | 1.78 |
| Ukraine, conventional | 2 | N/A | 1554 | 41.4 | 1119 | 1.74 |
| EU-27, conventional | 1 | N/A | 1300 | - | 1305 | 2.58 |

According to the Food and Agriculture Organization of the United Nations (FAO), sunflower oil was produced by 79 countries in 2014, though 53 of these contributed less than 0.5 % to global production^10^. Over 95% of production was from 24 countries, eight of which are represented by systems in this analysis. Interestingly, 28% of global sunflower oil supply in 2014 came from Ukraine, which is represented here by only two reports, whilst an additional 26% came from the Russian Federation, despite no relevant studies from this country having been identified in this analysis. It is likely that regional life cycle inventory reporting is impacted by country-specific political priorities. Where countries place a greater emphasis on agricultural sustainability and reducing GHG emissions, more research into this subject can reasonably be expected, which may reflect the disparities observed here. Since data on production in Ukraine and the Russian Federation, the two largest producers, were limited, we decided not to weight average data presented here by country-wide production in order to avoid false conclusions.

Studies included in the meta-analysis generally failed to provide inventory data on all five stages included within the system boundaries (Figure 1). A total of 31 out of the 32 included studies reported inventory data from the cultivation stage. Two studies reported data on seed drying. Eleven studies provided “distance to processing facility” data used to generate transport stage inventory data, processed here in terms of tonne kilometres (tkm). Eight studies reported data on oil extraction and refining. Finally, two studies reported inventory data on packaging. In order to normalise reported data items, whilst enabling missing values to be filled, all reports were consolidated into 22 systems, based on geographic location of each report and then by production system where relevant (Table 2). For each inventory data item, the mean average of values recorded across all reports in each system was calculated and used as the system inventory value.

After consolidation of reports into distinct systems, inventory data for most cultivation stage data items were available for each system. However, across all 22 systems and all 17 cultivation stage data items, a total of 84 out of 396 values were missing due to a lack of reporting. In these cases, the mean average value from all other studies for that data item was calculated and imputed. For the seed drying stage, only the “Italy, conventional” system was represented. Thus, data item values for all other systems were imputed, weighted by system yield. Ten systems had “distance to processing facility” data available. The mean average distance was calculated from these data and assigned to each of the other systems, prior to these data being used to calculate a system-yield weighted tkm value for each system. Processing stage inventory data was available in varying degrees for six out of the 22 systems. Once again, missing values were imputed based on system-yield weighted averages of available values for each data item. Finally, packaging inventory data was only available for two systems from two individual studies. Due to the lack of data on this production stage, this data was consolidated into a single set of data items, which was then yield-weighted and applied to all systems.

System-yield varied from 943 to 3287 kg ha^−1^ (Table 2). Interestingly, both highest and lowest yielding systems were from production in Portugal, the latter including irrigation. Meanwhile, oil recovery, reported for six of the 22 systems, ranged from 33% (w/w) for cold press systems in Italy, to 41.4% in Ukraine under conventional processing. It is worth noting that the literature search strategy employed in this study didn’t specifically seek to identify variation in seed or oil yield across production systems, hence this data can be considered only indicative.

### Organic production associated with lowest per hectare emissions, but conventional production can be more sustainable per kg oil produced

On a per hectare basis, total GHG emissions from sunflower oil production varied from 592 to 3822 kg CO_2_e ha^−1^ (Table 2). The lowest per hectare emissions were associated with production in Italy under organic cultivation, with oil extracted by cold-pressing. The highest per hectare emissions were associated with Iran, under conventional cultivation and oil extraction processes. Mean, systems-wide GHG emissions were 1396 kg CO_2_e ha^−1^, with a median of 1298 kg CO_2_e ha^−1^. Per kg processed, packaged oil produced, total GHG emissions varied from 1.08 to 4.19 kg CO_2_e kg oil^−1^ (Table 2; Figure 4). Expressed in these terms, the lowest total GHG emissions were associated with production in France, specifically with a legume included in the rotation and with a cover crop used in periods when land wasn’t cropped. Production in Iran was again associated with the highest GHG emissions. Mean emissions across all systems were 1.85 kg CO_2_e kg oil^−1^, with a median of 1.75 kg CO_2_e kg oil^−1^ (Figure 4).

**Figure 4.**
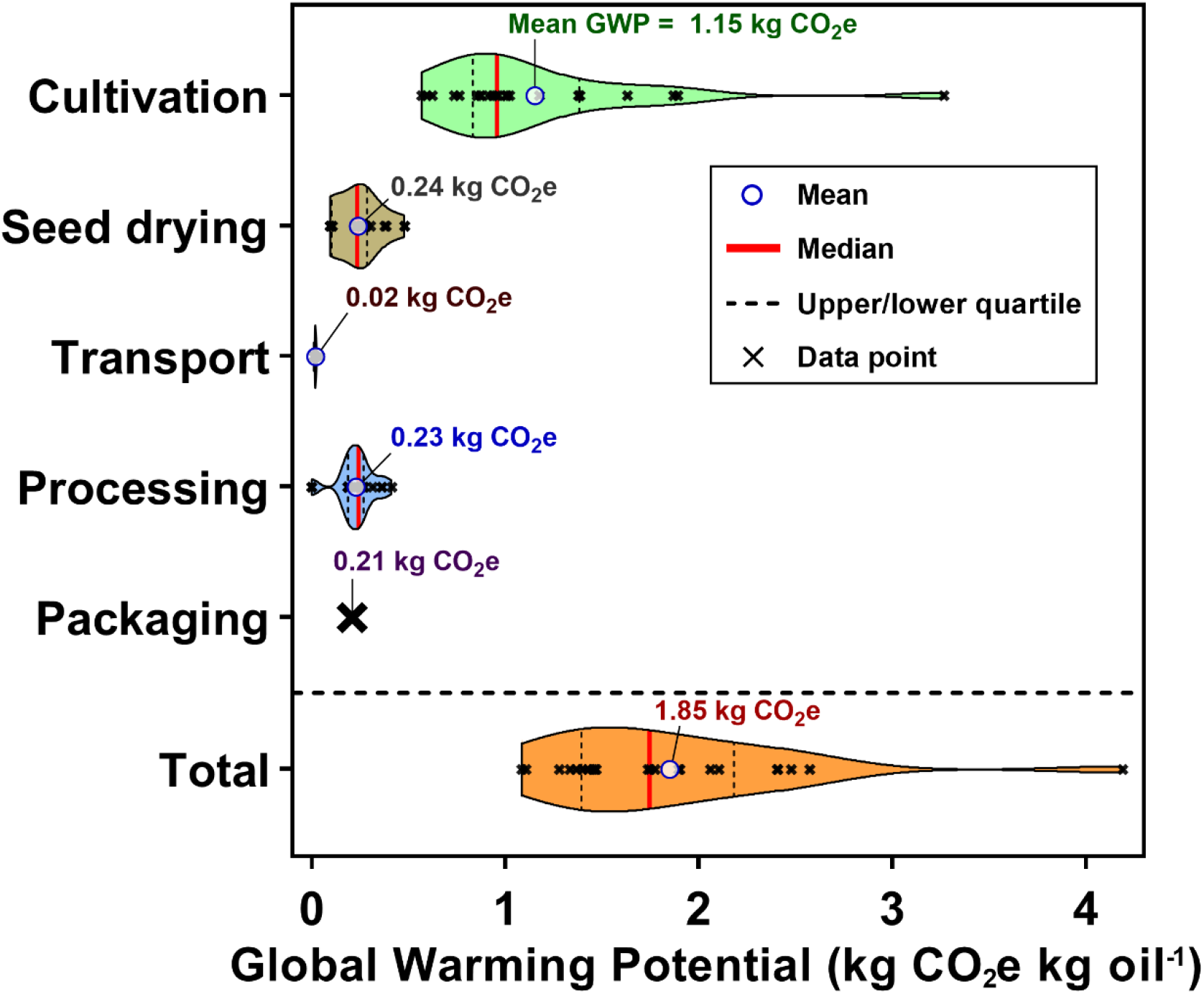
Life cycle greenhouse gas emissions across systems included in the life cycle meta-analysis. Emission sources grouped into five production stage categories with total life cycle emissions shown below. Violin plots represent full spread of data, with mean, median and upper and lower quartiles shown. Individual data points represented by crosses.

The majority of GHG emissions associated with sunflower oil production originated from cultivation, which contributed 62% of total emissions. Seed drying, processing and packaging contributed roughly the same as each other to total emissions; 13, 12 and 11%, respectively. Meanwhile, transport to the processing plant contributed only 1%. Variation existed within each production stage apart from packaging, for which only limited data was available for emissions calculations. Cultivation emissions varied almost 6-fold across systems, contributing from 46% (in Greece) to 79% (Italy, under conventional cultivation and cold press processing) to total GHG emissions. Transport emissions varied 4-fold, seed drying almost 5-fold, whilst processing emissions varied over 1800-fold, due to markedly fewer emissions originating from cold-press systems compared with conventional pressing systems. Despite large amounts of variation, processing only contributed up to a maximum of 22% of total GHG emission, as was the case for production in Greece (Figure 4).

### Diesel and nitrogen fertiliser use are main systems-wide contributors to GHG emissions

Diesel used during cultivation varied across systems from 50 to 135 kg ha^−1^, with a mean and median of 100 kg ha^−1^. Total synthetic nitrogen (as N) applied varied from 0 to 150 kg ha^−1^, with a mean of 55 kg ha^−1^ and a median of 54 kg ha^−1^. On average, cultivation diesel was the single largest emissions source across systems, contributing 32% to total GHG emissions. This was followed by N fertiliser use, which contributed a further 17% to total emissions on average across systems. Across the 22 consolidated systems under study here, two did not report use of any N fertiliser. These systems therefore reduce the average contribution of N fertiliser to total emissions when expressed in these terms.

Variation was observed in contribution of individual sources to total GHG emissions between production systems. For sunflower oil produced in Italy under conventional agronomic practice and cold-press processing, N fertiliser formed the single largest emissions source, contributing 43% to total emissions. Cultivation diesel contributed a further 28% to total GHG emissions in this system. Under conventional agronomic practice and oil processing in Italy, N fertiliser contributed 33% to total emissions, whilst cultivation diesel contributing a further 27%. Meanwhile, under organic production in Italy using cold-press processing, diesel contributed as much as 59% of total GHG emissions, due to fewer emissions originating from other sources. At the other end of the scale, diesel use in Iran contributed only 11%. This was despite roughly average diesel use in Iran compared to other systems; around 105 kg ha^−1^, compared to 99.8 kg ha^−1^ across all systems on average. The cause of apparently low contribution of diesel to total emissions in Iran is the large contribution of electricity for irrigation reported, which represents 42% of total system emissions. The latter parameter drove much of the high total GHG emissions of 4.19 kg CO_2_e kg oil^−1^, which reduced the contribution of diesel and other sources to total emissions proportionately. Within other stages of sunflower oil production, notable contributors to included steam production for pressing and processing of oil, which contributed 7.3% on average to total, per kg oil GHG emissions, and production of PET plastic used for packaging the finished product, which contributed a further 8.1% to total GHG emissions on average across systems.

### Yield is a significant driver of per hectare GHG emissions, whilst irrigation electricity and fertiliser use drive system differences in emissions per kg oil

Sunflower seed yield played a role in system differences in life cycle GHG emissions. On a per hectare basis, yield had a significant positive relationship with global warming potential (GWP; *r* = 0.45, *p* = 0.038; Figure 5a, 5b). This was due to a combination of a greater amount of processing inputs needed to process the greater volumes of seed produced, and to more intensive cultivation practices required to achieve a greater yield. On a per kg oil basis, this relationship was flipped, with a weak negative relationship observed between system seed yield and GWP (*r* = −0.28; Figure 5a, 5c), although this relationship was not statistically significant. It is worth noting that production in Iran was associated with a GWP of 4.19 kg CO_2_e kg oil^−1^, despite this system only producing a relatively average seed yield of 2339 kg ha^−1^. Hence, this system didn’t follow the general trend between yield and GWP (Figure 5b, 5c). Removing this system from the analysis resulted in a more strongly negative and statistically significant relationship (*r* = −0.57, *p* = 0.0067). However, results from production in Iran represent a genuine system, and thus are retained in all figures and tabular reporting here.

**Figure 5.**
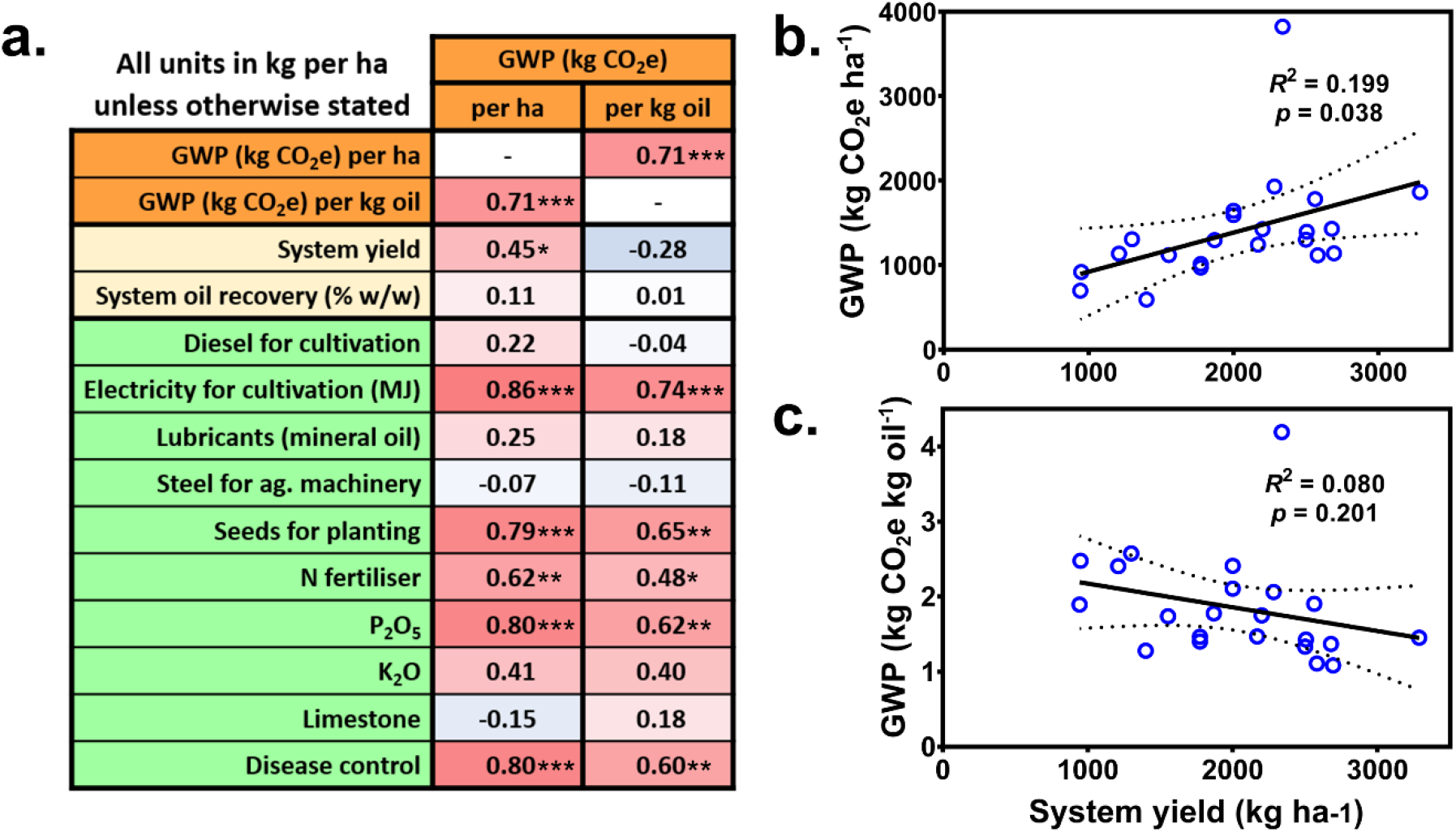
**a.** Correlations between system yield, oil recovery and cultivation stage emissions, and global warming potential (GWP) expressed per hectare (ha), and per kg refined, packaged oil. Cells are colour-coded with r values shown, where ‘more red’ represents a more strongly positive correlation and ‘more blue’ a negative correlation. Two-sided tests of correlation against zero significance levels shown where relevant (* = p < 0.05, ** = p < 0.01, *** = p < 0.001). **b/c.** Regression analysis between system yield and GWP expressed per ha (**b.**) and per kg packaged oil (**c.**). Regression line shown as solid line with 95 % confidence intervals shown as dotted lines.

As the single largest contributor to total emissions from production in Iran, electricity required for irrigation was also the input with the highest correlation with GWP expressed on both a per hectare and per kg oil basis (Figure 5a). Three out of the 22 systems reported here made use of irrigation during the cultivation stage, and two of these were associated with the first and second highest total GHG emissions. The first was associated with production Iran, and the second reflects production in the EU (EU-27) as a whole.

Unsurprisingly, as a large component of total GHG emissions, variation in N fertiliser used was positively correlated with GWP on both a per hectare (*r* = 0.62, *p* = 0.0021) and a per kg oil (*r* = 0.48, *p* = 0.025) reporting basis (Figure 5a). The use of phosphate fertiliser was also positively correlated with GWP both per hectare (*r* = 0.80, *p* < 0.001) and per kg oil (*r* = 0.62, *p* = 0.002), suggesting that this parameter plays a role in driving variation between systems GHG emissions estimates. However, much of this relationship is likely due to high phosphate fertiliser use reported for production in Iran, in which emissions are mostly driven by electricity used for irrigation. Thus, this positive relationship may be only indicative and not causal. Diesel use during cultivation, on the other hand, was not statistically correlated with GWP. This indicates that although diesel use makes up a large proportion of total GWP, there is limited variation in this parameter between systems and hence this doesn’t drive system differences.

### Additional spurious relationships identified with cultivation data items, but limited processing reporting limits further comparison

There were some perhaps more surprising positive correlations between GWP, and variation in both the quantity of seeds used for planting and disease control chemicals used (Figure 5a). This is despite seeds for planting only contributing 0.3% to total GHG emissions, and disease control chemicals contributing 3.0%. These relationships were likely driven instead by electricity used for irrigation, which was highly correlated with both seeds for planting (*r* = 0.81, *p* < 0.001) and disease control (*r* = 0.81, *p* < 0.001), and is a notable driver of total emissions in systems where it is present. In particular, relatively large inputs of both seeds and disease control were reported for production in Iran, which was associated with the highest GHG emissions among systems. After removing Iran from the analysis, the observed trends between these parameters disappear. Thus, the trends between seeds for cultivation and disease control chemicals used, and total GWP, can largely be considered spurious.

There were no statistically significant trends between emissions sources from any of the other sunflower oil production stages. This was due to limited reporting of seed drying, oil processing and packaging steps in the literature, leading to limited observable variation across systems for stages post-cultivation. For example, electricity used for processing was only reported for six out of 22 systems. However, there was considerable variation in emissions from electricity between these six systems; electricity contributed an average of 105 g CO_2_e per kg oil produced under conventional processing, but only 0.2 g CO_2_e per kg oil under cold-press processing. This was typical of differences observed across processing stage emissions. These trends are masked in the analysis by the systems for which processing inventory data is absent. In these cases, data values were inferred from the few studies for which data was available. Whilst methodologically crucial for estimating total life cycle GHG emissions, an unfortunate result of this is apparent low levels of variation across later production stages. It is clear that further reporting of inventory data for these stages across diverse production systems is required to allow proper comparison of associated systems-wide GHG emissions.

## Discussion

### Main emissions contributors indicate routes to more sustainable production

Variation in global sunflower oil GHG emissions was previously estimated in 2018 to between 2.2 and 4.9 kg CO_2_e per litre refined, packaged oil^6^. This is in agreement with results reported in the present study, which estimate GHG emissions between 1.1 and 4.2 kg CO_2_e per kg oil. The earlier estimate was representative of 519 farms, as opposed to 995 here, and was part of a larger study on environmental impacts across the entire food system. In contrast, the results reported here place stronger emphasis on variation in emissions associated with sunflower oil production across wide-ranging production systems. Our approach involved the re-analysis, using a single harmonised methodology, of primary data extracted from published studies, enabling a robust, cross-study re-analysis, incorporating a greater number of relevant reports, whilst allowing a detailed comparison of major emissions sources between systems. Such reporting will be highly useful for indicating routes to more sustainable production.

Three out of the 22 systems studied here made use of irrigation during sunflower cultivation. The electricity required for this was associated with particularly high levels of GHG emissions, most notably in Iran. According to the meta-analysis, over three time more irrigation electricity was consumed during cultivation in Iran compared to the system with the next highest irrigation use (Portugal, irrigated). The emissions associated with this were exacerbated by the relatively high carbon footprint of electricity production in Iran; 258 g CO_2_e per MJ, compared to an EU average of 132 g (IFEU calculations based on IEA 2012 data^27^, via BioGrace^24^). Electricity for irrigation was not the major emissions source in the other two systems that made use of it, (Portugal, irrigated; EU-27, conventional), but still formed a notable contributor. Diesel was a further major contributor to total GHG emissions. However, limited variation in diesel use across systems indicates a consistent requirement, making recommendations for increased sustainability through this input difficult.

Despite associated emissions being partially offset by a generally greater yield, N fertiliser was still strongly, positively correlated with GHG emissions per kg oil (Figure 5a). Two of the systems studied here used no N fertiliser (Italy, cold press, organic; Portugal, rainfed), and both were associated with lower than average per hectare emissions. However, both systems were relatively low yielding, which limited the per kg oil emissions savings. In contrast, production in Portugal using moderate N inputs (Portugal, irrigated) was associated with a high yield and lower than average GHG emissions per kg oil. Therefore, it seems that limiting N fertiliser use is not necessarily a suitable means to reduce the carbon footprint of sunflower oil production. Indeed, it was previously demonstrated that replacing all food production in England and Wales with organic produce would lead to greater net GHG emissions, due to lower yields requiring greater import of produce from other countries^28^. However, production in France with a legume in the rotation was associated with the lowest per kg oil emissions, despite very low levels of N having been applied. Lower N inputs were required due to the presence of residual soil N following legume cultivation^29^. Investment in production practices such as this, as well as work to understand the GHG response to varied N application rates, would go some way towards optimising N use for more sustainable production.

### Yield substantially influences GHG emissions and partially offsets impact of chemical inputs

This study identified a positive relationship between yield and per hectare global warming potential (GWP). Since the Green Revolution, crop productivity has been increasingly linked with the use of synthetic inputs, at high environmental cost^30^. Consistent with this, many of the systems that yielded higher in this study were associated with higher inputs of fertilisers and other chemicals. A higher yield also resulted in a greater quantity of seed requiring processing and packaging. Thus, the additional environmental burden associated with increased processing inputs further added to total GWP in this study when expressed on a per hectare basis. In contrast, an inverse relationship was observed when comparing yield with per kg oil GWP. This implies that increased yield led to production of a sufficient quantity of refined, packaged vegetable oil to partially offset the negative environmental effects that production inputs were associated with. In keeping with this finding, whilst positive correlations were identified between multiple agricultural inputs and GWP on both a per hectare and per kg oil basis, the relationship per kg oil was generally slightly weaker, and thus less associated with systems variation in total GWP (Figure 5a).

It is worth noting that as production of sunflower oil increases, so does the production of sunflower cake, a high protein meal used as animal feed. It is reasonable to assume that this may go some way towards offsetting alternative sources of animal protein and thus GHG emissions from alternative food systems. The implications of this are likely to be highly complex, given that a major source of high protein animal feed globally is soybean, which itself produces vegetable oil as a co-product of lower economic value^31^. It is therefore possible for an increase in sunflower cake production to decrease demand for and hence production of soybean, which could feedback an increase in demand for vegetable oil and thus sunflower production. However, the demand for animal feed is thought to be growing at a slightly faster rate than that of vegetable oil^13^. Thus, any increase in animal protein meal production from sunflower would likely fill new demand rather than reducing the need for alternative sources.

### GHG implications of meeting growing demand for vegetable oil with sunflower determined, laying the groundwork for future comparison

In this meta-analysis, we have evaluated the regional and systems variation in GHG emissions associated with sunflower oil production. However, this study has also highlighted limitations in available sunflower oil life cycle inventory reporting. Most notably, only two relevant reports were identified concerning sunflower production in Ukraine, despite this being the single largest producer at a country level^10^. Additionally, no reports were identified concerning production in the Russian Federation, which is the second largest producer^10^. Refining of systems-wide estimates of variation in GHG emissions from sunflower oil production would benefit from additional data concerning these regions. Nevertheless, the data presented here make significant progress towards harmonising GHG emissions estimates from 33% of the countries that contribute to the vast majority of worldwide sunflower oil production. These form a highly useful resource for informing the GHG implications of meeting additional demand for vegetable oil with sunflower. The findings produced here also lay the groundwork for comparison of sunflower with other major vegetable oil sources, such as palm, soybean and rapeseed, in order to identify the most environmentally sustainable system for global edible oil production.

## Author contributions

TDA performed systematic literature search, data extraction and analysis, and led the writing of the manuscript. DES and SJR contributed to project conception and oversaw project management and completion.

## Funding

TDA was funded by the Future Food Beacon of Excellence, University of Nottingham, UK. The authors declare that they have no competing interests.

## Supplementary material

Supplementary Table 1 – Literature search terms with number of results from each bibliographic database

Supplementary Table 2 – Data items collected, conversion factors used where relevant, and actions taken where values were missing from the consolidated inventory database

Supplementary Table 3 – List of literature from which inventory data was extracted for inclusion in the meta-analysis.

## Notes

### Competing Interest Statement

The authors have declared no competing interest.

